# HLA-G gene editing: a novel therapeutic alternative in cancer immunotherapy

**DOI:** 10.1101/2021.01.21.427294

**Authors:** María Belén Palma, Diana Tronik-Le Roux, Guadalupe Amín, Sheila Castañeda, Alan M. Möbbs, María Agustina Scarafia, Alejandro La Greca, Marina Daouya, Isabelle Poras, Ana María Inda, Lucía N. Moro, Edgardo D. Carosella, Marcela N. García, Santiago G. Miriuka

## Abstract

Cancer immunotherapies based mainly on the blockade of immune-checkpoint (IC) molecules by anti-IC antibodies offer new alternatives for treatment in oncological diseases. However, a considerable proportion of patients remain unresponsive to them. Hence, the development of novel clinical immunotherapeutic approaches and/or targets are crucial. In this context, targeting the immune-checkpoint HLA-G/ILT2/ILT4 has caused great interest since it is abnormally expressed in several malignancies generating a tolerogenic microenvironment. Here, we used CRISPR/Cas9 gene editing to block the HLA-G expression in two tumor cell lines expressing HLA-G, including a renal cell carcinoma (RCC7) and a choriocarcinoma (JEG-3). Different sgRNA/Cas9 plasmids targeting *HLA-G* exon 1 and 2 were transfected in both cell lines. Downregulation of HLA-G was reached to different degrees, including complete silencing. Most importantly, HLA-G – cells triggered a higher *in vitro* response of immune cells with respect to HLA-G + wild type cells. Altogether, we demonstrated for the first time the HLA-G downregulation through gene editing. We propose this approach as a first step to develop novel clinical immunotherapeutic approaches in cancer.

## Introduction

Cancer immunotherapy has improved outcomes of oncological treatments. Most of such breakthroughs are due to the discovery and therapeutic modulation of key immune-regulatory molecules (checkpoints) at the interface between tumor and immune cells ^1^. The interaction is now globally known as immune-checkpoints (IC), which have been broadly defined as cell-surface molecules that can transduce signals into effector cells to positively (stimulatory receptors) or negatively (inhibitory receptors) modulate signaling for preventing or promoting tumor cell survival, respectively^2^. IC blockade is now recognized as an effective therapy against some cancers.

The most extensively used in cancer immunotherapies are monoclonal antibodies directed to cytotoxic T-lymphocyte antigen 4 (CTLA-4) and Programmed Cell Death Protein 1 (PD-1)^3^. These therapies have been already extensively tested in clinical trials and have shown success in cancer treatment. However, the clinical effectiveness has been limited in some cases, possibly due to alternate pathways that are also critical in cancer. Moreover, the use of anti-CTLA-4 and/or anti-PD-1 antibodies is often associated with several adverse events such as immune-associated toxicity, treatment resistance, and autoimmune-like reactions, hence, the clinical benefit is limited to a fraction of patients^4–6^. Thus, the identification of new therapeutic targets or alternative therapies to improve patient survival and clinical outcomes is critical.

The interaction between HLA-G and its receptors ILT2 (LILRB1/CD85j) and ILT4 (LILRB2/CD85d) is an IC that has generated a great interest in the past years as putative immunotherapy target^7,8^. HLA-G is a non-classical MHC class I molecule that was originally described in trophoblast cells at the maternal-fetal interface where it plays a critical role in protecting fetal allograft tissue from maternal immune rejection^9^. Its primary transcript undergoes alternative splicing, producing at least seven mRNAs encoding four membrane-bound (HLA-G1 to HLA-G4) and three soluble (HLA-G5 to HLA-G7) protein isoforms, all beginning at the same translation start site situated in exon 2^10^. Interestingly, no distinct functional roles have yet been described for these isoforms. Recently, two novel isoforms were described^11^. The first one lacks exon 3, resulting in a transcript that encodes an isoform lacking the α1 domain. The other isoform is transcribed from a supplementary exon previously unknown, which contains an upstream ATG. The translation from this ATG located in exon 1 can generate a 5 aa-extended N-terminal protein. Moreover, the presence of this exon may alter RNA stability and translation by modifying the binding of regulatory proteins and/or micro-RNAs.

HLA-G has a broad immunoregulatory function that affects both innate and adaptive immunity. Through its interaction with the inhibitory receptors ILT2 and ILT4, both mainly expressed by immune cells, HLA-G exerts different immune regulatory functions, including inhibition of the cytolytic function of NK cells, the antigen-specific cytolytic function of cytotoxic T cells, the alloproliferative response of CD4+ T cells and the maturation of dendritic cells^8,12^. Moreover, unlike other IC, HLA-G has a restricted expression in normal tissues, such as in thymus, cornea, some activated monocytes, and erythroid and endothelial precursors^13–15^. However, its expression can be ectopically induced under malignant cell transformation in tumor and/or in tumor-infiltrating immune cells^16,17^. Thus, HLA-G is a promising target for new immunotherapies with relatively low chances of significant side effects.

There is a growing amount of promising preclinical data showing that Clustered Regularly Interspaced Short Palindromic Repeats (CRISPR)/CRISPR associated nuclease 9 (Cas9) constitutes a powerful gene-editing tool to specifically target cancer cells and suppress tumor growth^18–20^. CRISPR/Cas9 has revolutionized genetic engineering and it is emerging as a robust alternative strategy for current cell-based immunotherapy that will minimize potential side effects caused by antibody blockade therapies. Gene editing allows the generation of site-specific modifications at a very specific point of the gene code, inactivating it by the incorporation of insertions or deletions (InDels) in its sequence. Considering some limitations that the use of anti-IC antibodies have when interfering with IC, the development of alternatives such as CRISPR/Cas9 gene editing is a promising strategy.

Here, we used CRISPR/Cas9 gene editing to disrupt *HLA-G* gene expression in tumor cell lines. We generated several clonal cell lines with different HLA-G – degrees of downregulation, which constitutes a proof-of-concept study regarding the feasibility of knocking down the HLA-G gene in tumor cells. The final goal is to interfere of the HLA-G/ILT-2 or ILT-4 immunological synapse and restore the host immune capacity to attack the cancer cells. We propose this approach as a first step to develop a new alternative for cancer therapy.

## Results

### 1. Transfection of CRISPR/Cas9 system in the RCC7/HLA-G1 cell line

The RCC7 cell line previously transduced with a lentivirus containing HLA-G1 cDNA, was transfected with pSpCas9 vector cloned with the 2A-sgRNA that target exon 2 region (Fig. 1). One hundred clonal cell lines were obtained following cell sorting with FACSAria III. The HLA-G protein levels of several selected clones were compared with the wild type RCC7/HLA-G1 by Western blot (WB). We observed a significant downregulation of HLA-G protein expression for most of the transfected clones with the 2A-sgRNA (Fig. 2A). We then selected three representative clonal cell lines (AA4, AA8 and EE2) to further analyze the modifications that had occurred at the mRNA level. To this end, three different combinations of primers that target exon 1, exon 2 and exon 3-4 were used. The RT-PCR results showed the expected amplification fragments with the three pairs of primers for the wild type RCC7/HLA-G1 cell line. In the case of the selected edited clones, no amplification could be obtained by using primers complementary to exons 1 or 2 whereas an expected fragment was obtained with the primers located in exons 3-4 (257F/526R), far downstream of the target sequence of the 2A-sgRNA (Fig. 2B). This demonstrates that a sequence edition occurred adjacent to the 2A-sgRNA target site.

**Figure 1:**
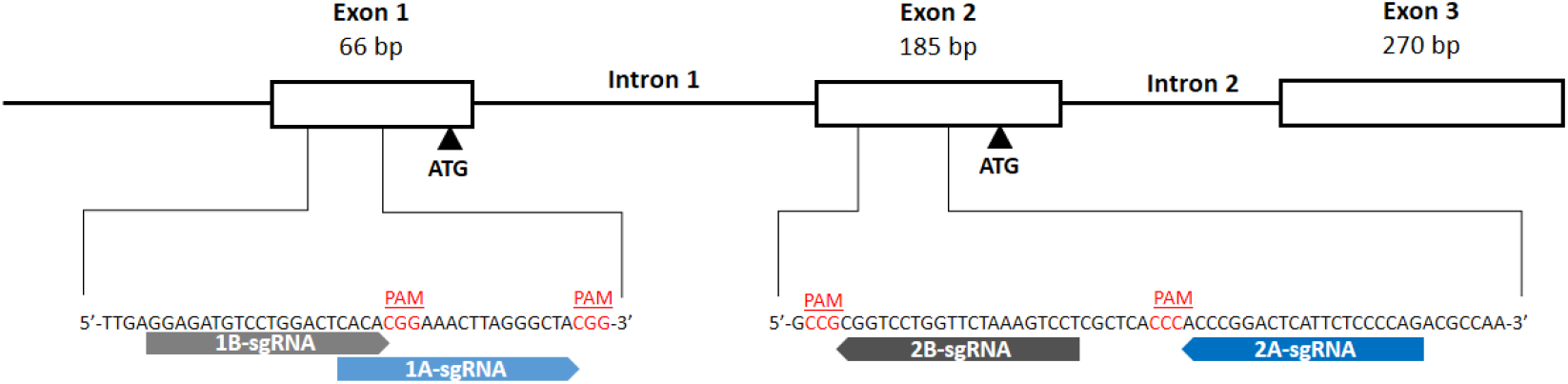
Schematic representation of the first 3 exons of *HLA-G* gene and the 4 designed sgRNAs. The white boxes and lines represent exons and introns, respectively. The sequence below represents part of exon 1 and 2 containing Cas9/sgRNA target sites for 1A-, 1B-, 2A- and 2B-sgRNAs. Protospacer-adjacent motif (PAM) is labeled in red. Black arrows represent the two transcription start sites.

**Figure 2:**
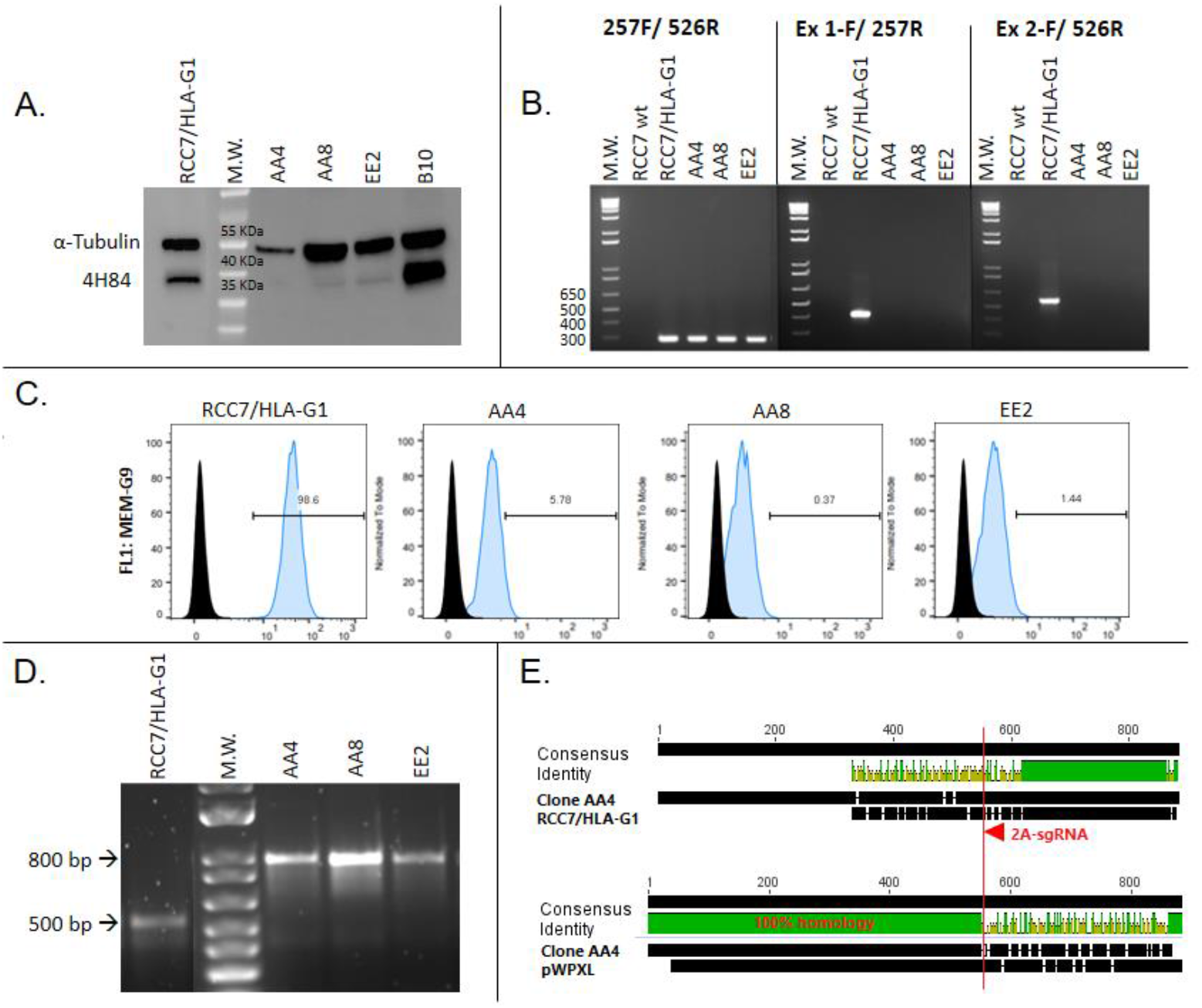
Analysis of RCC7/HLA-G1 edited cells by CRISPR/Cas9: **A.** Western Blot analysis of HLA-G expression in 2A-sgRNA clonal cell lines (right). Wild type RCC7/HLA-G1 control is shown on the left. **B.** RT-PCR to analyse mRNA expression. **C.** Flow cytometry analysis of HLA-G protein expression. **D.** Genomic DNA amplified by PCR. **E.** Sequence alignment of wild type RCC7/HLA-G1 *vs*. clone AA4 and pWXPL-lentivirus vector. The red arrow represents the 2A-sgRNA target site and the red line shows the cut site of Cas9 protein. http://www.geneious.com.

To more precisely quantify the HLA-G downregulation, we analyzed the three selected clones by flow cytometry (FC). The results showed HLA-G protein downregulation of 94.2%, 99.6% and 98.5% for each clone respectively (Fig. 2C). Then, to identify the genomic modifications occurring after the transfection of the sgRNA, the gDNA-edited region was amplified by PCR using primers CRR1F/257R. The results revealed an 800bp amplicon after PCR amplification, which is 300 bp longer than the expected fragment of 500bp (Fig. 2D). To determine the origin of this 300 bp-insertion that occurred in the 2A-sgRNA clonal cell lines, the fragments were sequenced. The alignment between the wild type RCC7/HLA-G1 and the edited clonal cell lines sequences revealed that the 300 bp insertion corresponded to a DNA fragment derived from the pWXPL-lentivirus vector into which the HLA-G DNA was cloned (Fig. 2E).

Overall, these results demonstrate that the designed sgRNA was suitable to achieve the downregulation of artificially expressed HLA-G, although it could be partly explained by the usage of the previous lentiviral backbone gene as a scaffold to introduce a genomic insertion.

### 2. Transfection of CRISPR/Cas9 system in JEG-3 cell line

Considering the previous interference of the lentivirus vector in where the HLA-G1 cDNA was cloned, we applied the gene editing strategy to the JEG-3 cell line that naturally expresses HLA-G. Also, to have a better insight into the regulation of the *HLA-G* gene, we designed three supplementary sgRNAs, named 1A-, 1B- and 2B-sgRNAs, that target the two reported translation start sites. According to the Ensembl Genome Browser (http://www.ensembl.org/index.html), the *HLA-G* gene has 8 exons (Human GRCh38.p13). The most important translation start codon is present in exon 2. However, there is another ATG in exon 1 that can be used as initial start codon when a 106 bp deletion occurs between exon 1 and 2^11^. The Figure 1 shows the four sgRNAs designed, 1A- and 1B-that targeting upstream exon 1 ATG and 2A- and 2B-that target upstream of the ATG situated in exon 2. The JEG-3 cell line was transfected with pSpCas9 vector cloned with each sgRNA and analyzed as follows.

First, HLA-G expression was measured by FC on JEG-3 edited cells. The results revealed that using any of the sgRNAs (1A, 1B, 2A or 2B), the cell membrane HLA-G expression was downregulated. Compared with the wild type JEG-3, the proportion of HLA-G reduction with each sgRNA was 72%, 35%, 80% and 62% for 1A, 1B, 2A and 2B-sgRNAs, respectively (Fig. 3A, B, and D). Moreover, transcriptomic analysis by RT-qPCR with primers located 3’ of the ATG start site (257F/526R) also confirmed these results, with a reduction of *HLA-G* mRNA of 60%, 40%, 65% and 55% for 1A, 1B, 2A and 2B-sgRNAs, respectively (Fig. 3A and B; data were normalized to the unedited cells, wt JEG-3).

**Figure 3:**
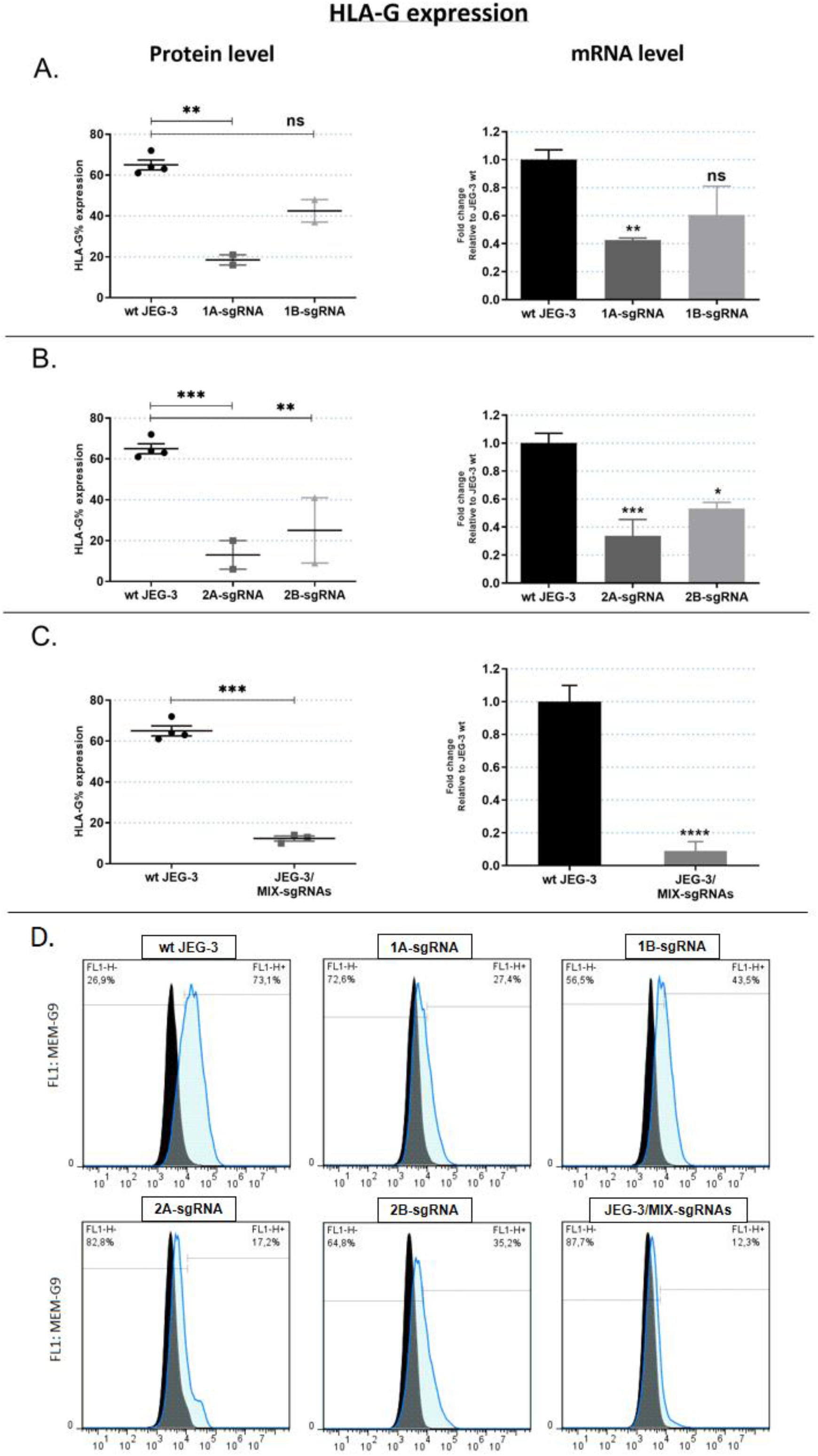
Analysis of JEG-3 edited cells by CRISPR/Cas9: **A.** HLA-G expression in JEG-3 edited cells in exon 1 with 1A- and 1B-sgRNAs. **B.** HLA-G expression in JEG-3 edited cells in exon 2 with 2A- and 2B-sgRNAs. **C.** HLA-G expression in JEG-3 edited cells using the four sgRNAs (1A-, 1B-, 2A- and 2B-sgRNAs, named MIX-sgRNAs). On the left the HLA-G protein expression was measured by FC. On the right the *HLA-G* mRNA expression was measured by RT-qPCR. No significant: ns. Significant differences are shown with * (p<0.05). **D.** Representative histograms of HLA-G measure by flow cytometry in wild type JEG-3 cells (wt) and cells edited in exon 1 (1A- and 1B-sgRNAs), in exon 2 (2A- and 2B-sgRNAs), or in both exons (JEG-3/MIX-sgRNAs). Isotype control is shown in black.

Finally, the genomic *HLA-G* sequence was analyzed in JEG-3 edited cells with each sgRNA to determine which editions occurred. The PCRs were performed using the CRD1 F/R oligonucleotides to amplify the exon 1 genome region, and the CRD2 F/R to amplify the exon 2 genome region. Using Synthego’s ICE tool, the percentage of edition was determined by comparing with the wild type sequence. We observed that effectively the genome was edited in all cases. Different InDels occurred with each sgRNA, with an accumulative genome modification rate of 63%, 88%, 79% and 71% for 1A, 1B, 2A and 2B-sgRNAs, respectively (Fig. 4A and B).

**Figure 4:**
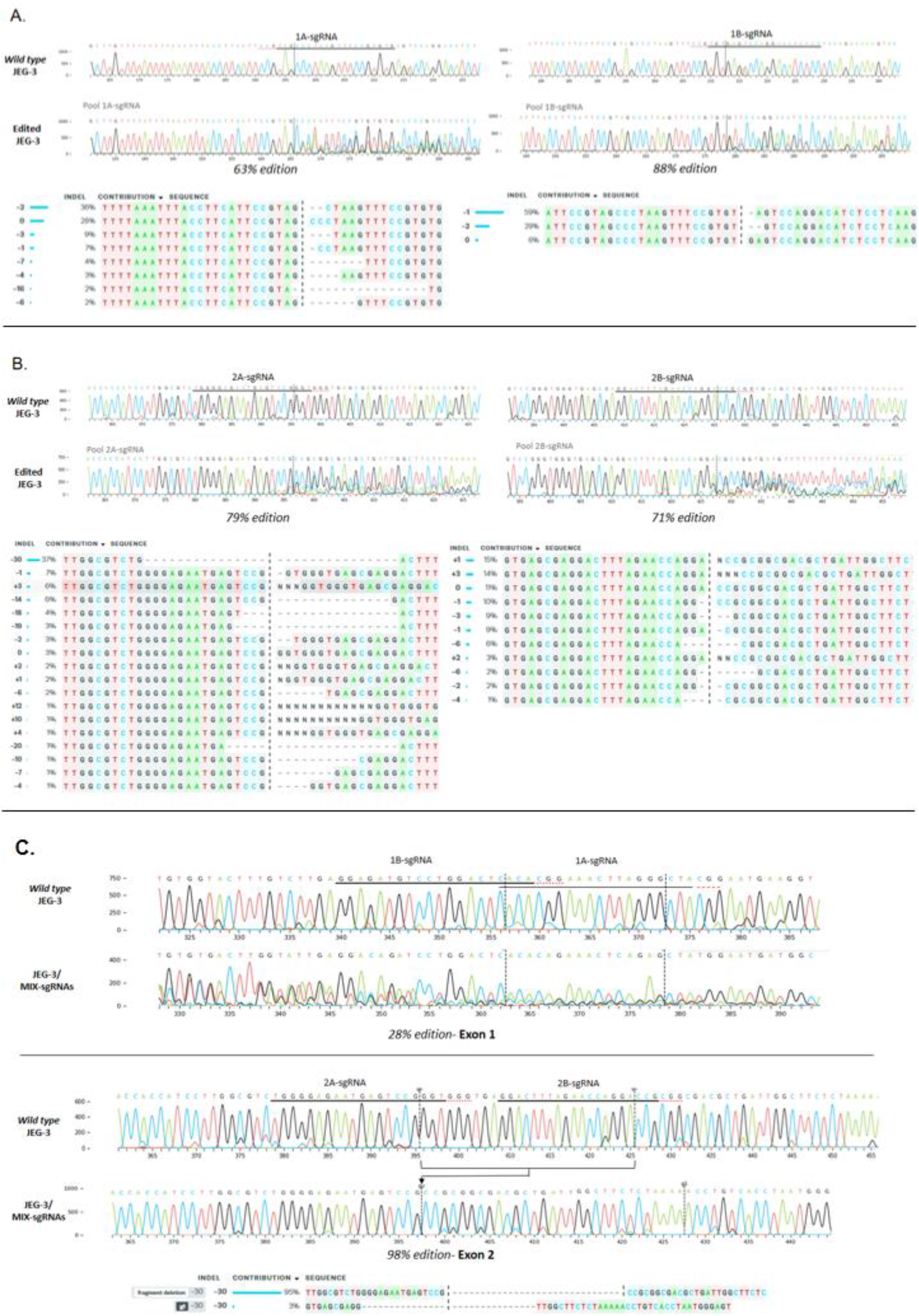
Sequencing analysis. Histograms with nucleotide sequence data of cell pools edited by CRISPR system. Different edited genotypes predicted by InDels analysis (https://www.ice.synthego.com/) and percentage of edition for each condition: **A.** JEG-3 cell pools edited with 1A- and 1B-sgRNAs, in exon 1 region. **B.** JEG-3 cell pools edited with 2A- and 2B-sgRNAs, in exon 2 region. **C.** JEG-3 cell pools edited with all sgRNAs (1A, 1B, 2A and 2B, named JEG-3/MIX-sgRNAs), in exon 1 and 2 simultaneously.

### 3. Transfection of CRISPR/Cas9 system with 4 sgRNAs simultaneously in JEG-3 cell line

The results mentioned above showed that the HLA-G expression was reduced, but not completely turned off when each sgRNA was used separately. To achieve a complete HLA-G-knockout cell line, JEG-3 cells were transfected with the 4 sgRNAs simultaneously (JEG-3/MIX-sgRNAs). The *HLA-G* mRNA (measured by RT-qPCR) and the protein expression (measured by FC) were 90% reduced in JEG-3/MIX-sgRNAs with respect to the wild type JEG-3 cells (Fig. 3C and D). Consistent with a completely disrupted HLA-G expression, both at transcriptomic and proteomic levels, results of the DNA sequencing showed that effectively in 98% of cases the cells were edited. Indeed, a deletion of 30 bp in exon 2 was observed between 2A and 2B-sgRNAs target sites (Fig. 4C). This genome modification is concordant with the significant reduction of HLA-G expression.

### 4. Analysis of NK degranulation in co-culture with HLA-G wt and HLA-G – JEG-3 cells

One of the important HLA-G functions is the inhibition of NK cell degranulation. This process can be measured by detecting the Lysosome-associated membrane protein-1 (LAMP-1 or CD107a) at the cell surface. In fact, as a consequence of the degranulation process, the outer membrane of the granules merges with the NK cell plasma membrane, leading to surface exposure of CD107a molecules. The HLA-G inhibitory function over NK cell degranulation was then determined after its stimulation and the co-culture with HLA-G wt (wt JEG-3) and HLA-G - (JEG-3/MIX-sgRNAs) cells. First, FC analysis determined the NK identity by anti-CD45/PE, anti-CD56/BB515 and anti-CD3/PE antibodies. The 97.1% of these cells were CD45 (+) and CD56 (+), and more than 60% of the CD56 (+) cells were CD3 (-), corresponding to NK lymphocytes. The CD56 (+) and CD3 (+) population corresponded to ɣ δ T lymphocytes^28^ (Fig. 5A). Then, we measured cell surface CD107a present in the NK cell population. Two control conditions were performed: a negative control including NKs without co-culture with target cells nor stimulation cocktail to analyze basal degranulation, and a positive control including NKs without target cells but with stimulation cocktail to activate the spontaneous degranulation process (Fig. 5B). NKs co-cultured with HLA-G – (JEG-3/MIX-sgRNAs) cells expressed 19.7% of CD107a compared to 14.9% when NKs were co-cultured with HLA-G wt (wt JEG-3) cells (p<0.01) (Fig. 5C), implying an increased NK degranulation when NKs were co-cultured with HLA-G - edited cells. These percentages are consistent with previous published results^29^.

**Figure 5:**
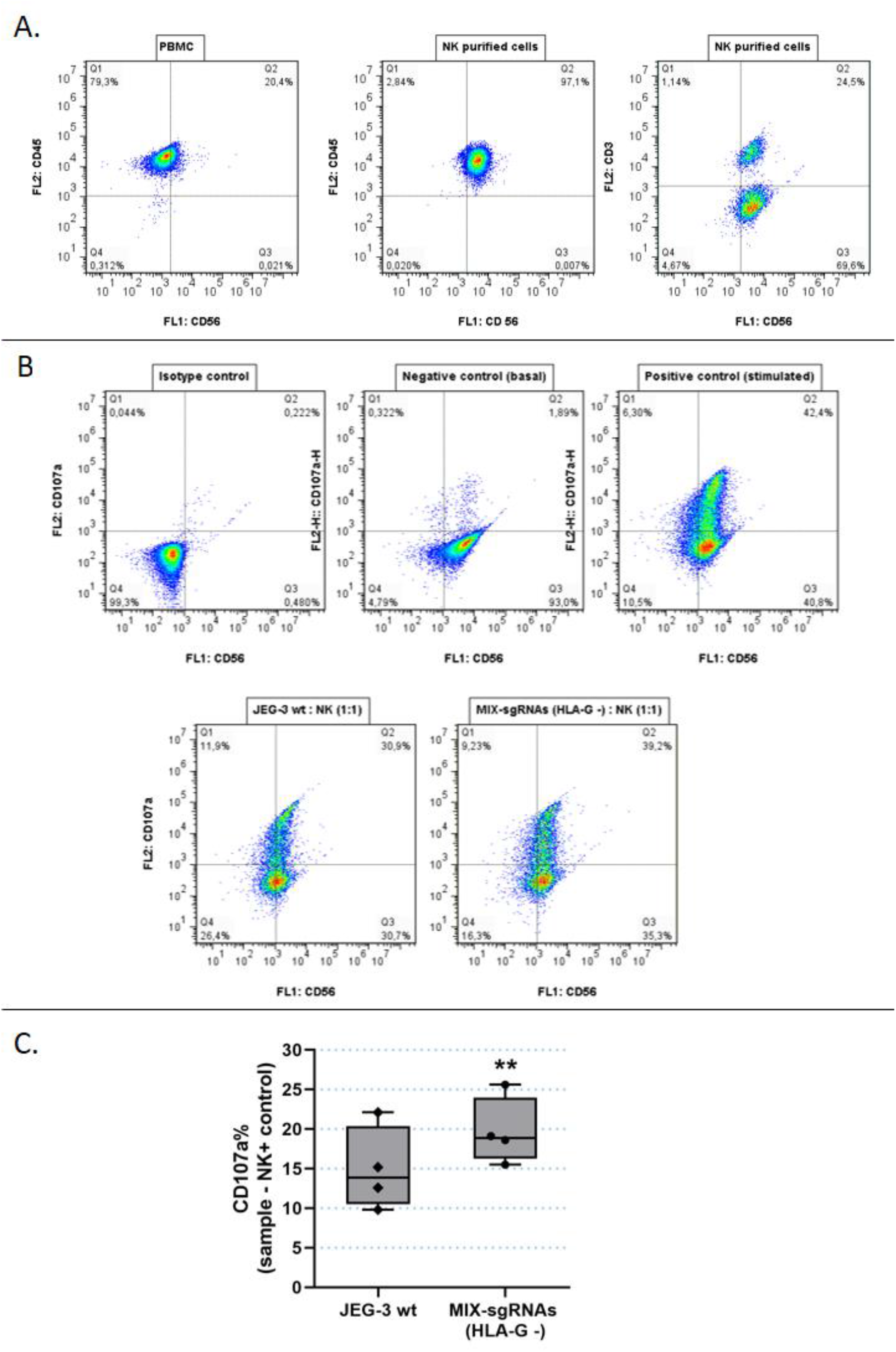
Degranulation assay, functional analysis of HLA-G wt and HLA-G - JEG-3 cells. **A.** Determination of NK cells purification. NK cells were stained with anti-CD45/PE, anti-CD56/BB515 and anti-CD3/PE and compared with PBMC. **B.** As a representative assay, NKs were labelled with CD107a/PE and CD56-BB515. The conditions were the following: Isotype control (NK without Ab), negative control (NK basal degranulation), positive control (stimulated NK cells) and NKs co-cultured with wt JEG-3 or with JEG-3/MIX-sgRNAs (HLA-G -). **C.** Box plots show the percentage of CD107a+ NK cells when co-cultured with wt JEG-3 (HLA-G wt) or with JEG-3/MIX-sgRNAs (HLA-G -). Significant differences are shown with * (p<0.05) (n= 4 experiments).

## Discussion

Immunotherapy has recently emerged as a viable and attractive treatment option for many cancer patients. In particular, monoclonal antibody-based IC blockade therapies that enhance the function of anti-tumor T lymphocytes have been particularly promising, and many therapies have been approved in several cancer types, such as renal cell, melanoma and lung cancer^30–36^. However, clinical trials have shown limited efficacy, a considerable proportion of patients did not respond to the treatment. Adding new therapeutic targets is then warranted.

Over the last decades, aberrant HLA-G expression has been found in numerous types of cancer, which has been associated with an advanced tumor stage, aggressive transformation and poor disease prognosis^37^. Furthermore, the low or null HLA-G expression in normal tissues makes it an interesting therapeutic target. Therefore, it has been proposed that the HLA-G blockade could be beneficial in any neoplasia that expresses HLA-G as an evasion mechanism of immune surveillance^38^. In this way, we proposed the HLA-G blockade by CRISPR/Cas9 gene editing, an effective tool to make genomic engineering manipulations, as a potential alternative therapy. This strategy could lead to re-activation of the host immune system to attack tumor cells. In this paper, we have used sgRNAs that target two genomic regions expected to affect the HLA-G translation in two different tumor cell lines.

*HLA-G* gene edition in RCC7/HLA-G1 cell line achieved a total protein silencing in all the clonal cell lines analyzed by WB and FC when the 2A-sgRNA was used. Also, when using specific oligonucleotides against exon 1 and 2 we did not obtain amplification by RT-PCR in the clones, which indicates that these regions were edited. The sequencing results corroborated that all clones were edited, and showed that a fragment of 300 bp was introduced into the genome. This insertion matched the sequence with the pWXPL-lentivirus vector. A possible explanation is that when the Cas9 endonuclease generated a double break in the DNA, it was probably repaired by homology-directed repair (HDR) using a second molecule of pWXPL as a template, also integrated into the genome. In any case, we confirm that the sgRNA was able to target this specific *HLA-G* genomic region.

The second edited tumor cell line was JEG-3, which naturally expresses high levels of HLA-G. At this time, we used four different sgRNAs (1A, 1B, 2A and 2B) to increase chances of disrupting HLA-G expression. The results showed that all the conditions were able to decrease HLA-G expression, though 2A and 2B-sgRNAs were more effective than 1A and 1B-sgRNAs. All the designed sgRNAs were able to recognize and edit the *HLA-G* genome, however, a single sgRNA was not enough to generate a complete HLA-G knockout cell population. This could be the result of editing only one of the translation initiation sites (exon 1 or 2), but not both of them. Hence, all sgRNAs were transfected simultaneously in order to increase the efficiency of the knockdown. We found that under this condition an almost total HLA-G silencing was achieved. Therefore, it was then demonstrated that the two ATG regions are perhaps similarly important in the translation process^39^. Finally, we showed that the HLA-G function was affected after gene editing measuring its influence over the NK degranulation process. Co-culture of NK with edited JEG-3/MIX-sgRNAs cells demonstrated that when the HLA-G expression disappears from the tumor cell surface, the NK cells partially recover their degranulation activity.

After gene editing, both *HLA-G* mRNA and protein decreased consistently. According to the theoretical framework of gene editing, in most cases, mRNA expression is normal, whereas protein expression is disrupted. This occurs when there are small variations, only InDels of few base pairs, that do not alter the mRNA expression but generates a frameshift mutation, and the protein will not be functional or is not translated. However, the Sanger-sequencing results showed an important edition in our edited cells, with InDels of several nucleotides, as shown in Figure 4. Based on InDels analysis (ICE, Synthego) of the cell pools, the edition average efficiencies were 63%, 88%, 79% and 71% for 1A, 1B, 2A and 2B-sgRNAs, respectively, and 98% with all four sgRNAs. These acquired mutations in the genome could harbor premature termination codons and mRNA could be degraded by Nonsense-mediated mRNA decay pathway^40,41^. Alternatively, these mutations could modify the core promoter and RNA polymerase could be inactive^42^. Hence, we found a consistency between higher editing rates and lower mRNA and protein expression.

Actually, gene editing is proposed in many cancer treatments^43,44^. One promising area in immunotherapy using CRISPR/Cas9 is its application on genetically engineered allogeneic T cells, known as chimeric antigen receptor (CAR) T cells. In several studies, CAR-T cells derived from healthy donors were edited to silence or disrupt both TCRs and HLA molecules to administrate in an allogeneic manner in oncological patients^18,45^. This allows the targeting of tumor-associated antigens and could enhance the therapy response by activation of T cells without host rejection. Another strategy to enhance the CAR-T cell therapy is to destroy the PD-1 expression by CRISPR/Cas9 system because this inhibitory signal generates T cell exhausted, and its blocking could be an improvement in the antitumor efficacy and clinical outcome^20,46^.

Gene editing is currently on the way to be applied in many different diseases and has an enormous potential^47^. However, there are still certain challenges that need to be overcome for safe and effective use of CRISPR/Cas technology in clinical gene therapy applications, such as delivery vehicles specific on target tissue, immunogenicity and DNA damage response, among others. In the future, to overcome these obstacles, we propose to deliver the CRISPR therapeutics into the human body using adeno-associated virus (AAV) vectors^48–50^. AAV is safe, capable of delivering the CRISPR/Cas system to various tissues and cell types, and only mildly immunogenic within a wide range of doses. Furthermore, the vector largely remains episomal inside host cells, it is stabilized through concatemerization and circularization to mediate long-term transgene expression in post-mitotic cells, leading to durable therapeutic efficacy.

In summary, we demonstrated for the first time that it is possible to block HLA-G expression in two different tumor cell lines through gene editing leading to its downregulation, with a concomitant effect in immune cell activation. This approach would reactivate the host immune system and help to eliminate tumor cells, thus proposing a novel immunotherapy.

## Methods

### 1. Cell culture

Two different cell lines with high levels of HLA-G were used: a renal cell carcinoma cell line (RCC7), derived from a clear cell renal cell carcinoma patient (kindly provided by Anne Caignard^21^), and a choriocarcinoma cell line [JEG-3, generously provided by Instituto de Fisicoquímica Biológica y Química, Universidad de Bioquímica y Farmacia (UBA-CONICET), Buenos Aires, Argentina]. The RCC7 line does not express HLA-G, but it was previously transduced with lentivirus containing HLA-G1 cDNA, generating a stable cell line expressing high levels of HLA-G1 (RCC7/HLA-G1)^22^. Instead, JEG-3 cells expresses HLA-G (see Supplementary Information Fig. S1).

Both cell lines were cultured *in vitro* in DMEM (Gibco) supplemented with 10% foetal bovine serum (Gibco) and 1% penicillin/streptomycin (Gibco) in a 5% CO2, humidified atmosphere at 37°C. Cells were regularly dissociated using Trypsine-EDTA 0.25% (Gibco).

### 2. Design and preparation of sgRNA vectors

Four sgRNAs were designed: two sgRNAs upstream exon 1 ATG (named 1A- and 1B-sgRNAs), and two sgRNAs upstream of the ATG situated in exon 2 (named 2A- and 2B-sgRNAs). All the sgRNAs were designed using Benchling Life Sciences R&D Cloud Software (https://benchling.com/) and cloned into the pSpCas9(BB)puroV2.0 vector (Addgene #62988), which expresses both Cas9 and puromycin resistance genes^23,24^. The sgRNA sequences are listed in Table 1 and the schematic representation of sgRNAs design is shown in Figure 1.

**Table 1:**
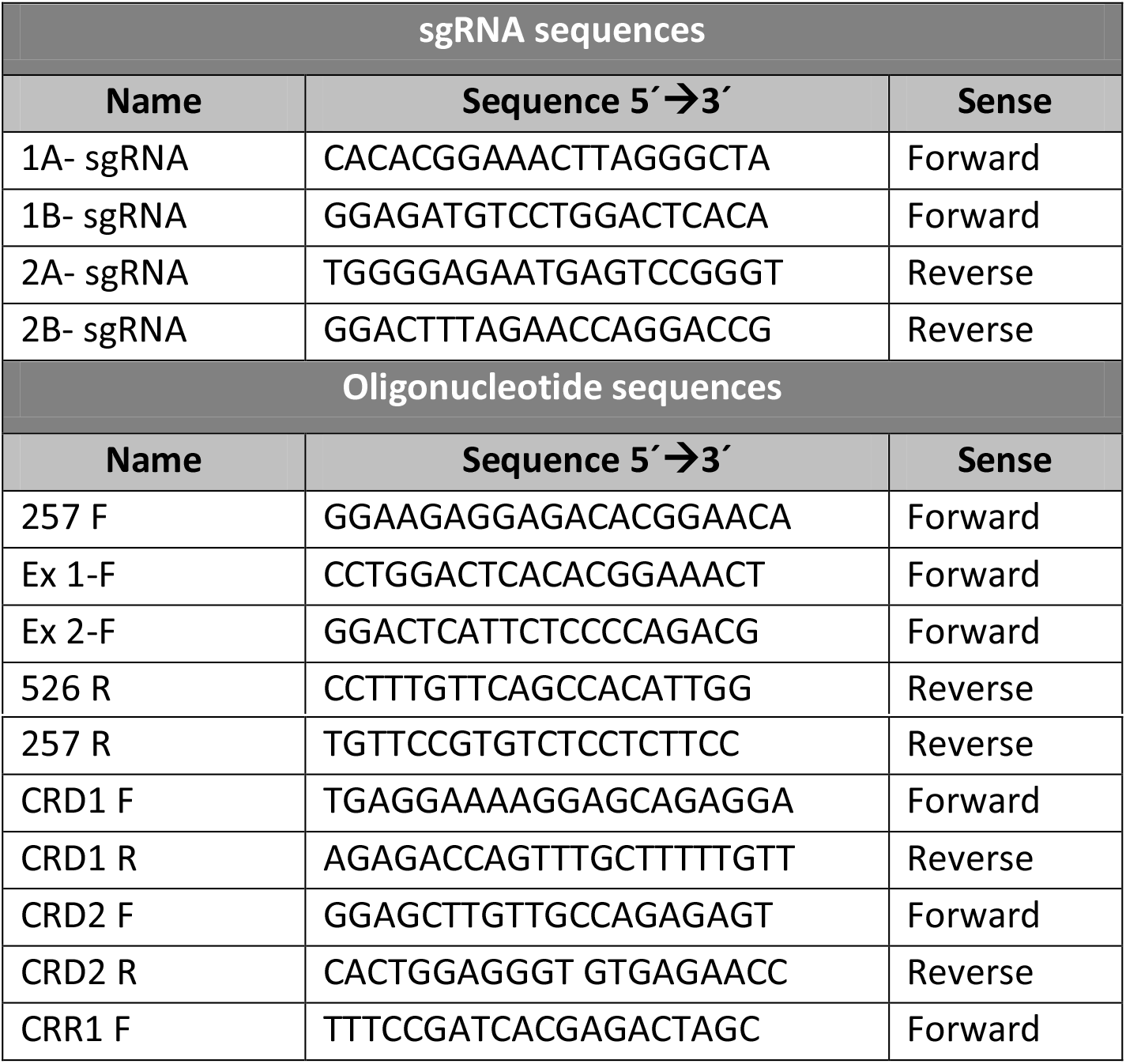
sgRNA sequences used for gene editing. Oligonucleotide sequences used in RT-PCR, RT-qPCR and gDNA amplification.

### 3. CRISPR/Cas9 vector construction and transfection

Transfection of CRISPR/Cas9 constructs was performed using X-tremeGENE 9 DNA Transfection Reagent (Roche), with 4.0 μg plasmid per 200.000 cells/well in a six-well plates, according to the manufacturer’s instructions. The RCC7/HLA-G1 cell line was transfected with 2A-sgRNA plasmid. The JEG-3 was transfected with 1A-, 1B-, 2A- and 2B-sgRNA plasmids separately, or with the four plasmids transfected together. As a control, we also transfected the cells with a GFP expressing vector (pEGFP-N1, Addgene #6085-1). Fluorescence images were captured with a Nikon Eclipse TE2000 inverted microscope (Nikon, Melville, NY, USA). Transfected cells were selected after 48 hs by adding 1.75 μg/ml of puromycin (Invivo Gene) and further cultured for another 48 hs.

For RCC7/HLA-G1 resistant cells, the clonal cell lines were obtained by sorting the cells with BD FACSAria III (BD Biosciences-US).

### 4. RNA Extraction, cDNA synthesis and real-time RT-qPCR

RNA extraction from puromycin resistant cells (RCC7/HLA-G1 and JEG-3) was performed with TRIzol Reagent (Invitrogen). For cDNA synthesis, 500-1000 ng of the total RNA was retro-transcribed with MMLV reverse transcriptase (Promega), according to manufacturer’s instructions. For RT-qPCR, cDNA samples were diluted 5-fold and it was performed with StepOne Plus Real Time PCR System (Applied Biosystems). The FastStart Universal SYBR Green Master Mix (Roche) was used for all reactions. Primers efficiency and initial molecule (N0) values were determined by LinReg software 3.0, and gene expression was normalized to *RPL7* housekeeping gene, for each condition. All the oligonucleotide sequences are listed in Table 1.

### 5. Western blot

WB analysis was performed to assess the expression of the HLA-G protein in RCC7/HLA-G1 after transfection of the CRISPR/Cas9 system by using the 4H84 mAb (Exbio) at a 1:1000 dilution. Peroxidase conjugated sheep anti mouse IgG Ab (Sigma) at a 1:1000 dilution was used as secondary antibody, as previously described^25^.

### 6. Flow cytometry

To determine the HLA-G silencing in JEG-3 cell line by CRISPR/Cas9, the protein expression was analyzed by FC. JEG-3 cells transfected with each sgRNA individually or all sgRNAs together, were dissociated and stained with a primary antibody anti-HLA-G conjugated with FITC (Invitrogen), in a 1:50 dilution for 30 min at room temperature. FC analyzes were performed in a BD Accuri cytometer. Data was analyzed with FlowJo Software.

### 7. Genomic sequence analysis

Genomic DNA extraction was performed using lysis buffer (10 mM Tris-HCl pH 8.3, 50 mM KCl, 2 mM MgCl2, 0.001% gelatine, 0.5% NP-40, 0.5 % Tween-20) and 0.05 mg/ml of proteinase K (Invitrogen). Following that, the gDNA was purified and stored at −20°C. Oligonucleotide sequences used to amplify the modified genome region are listed in Table 1.

PCR was performed using Easy Taq DNA Polymerase (Transgen Biotech). The desired PCR fragments were isolated from a 1% agarose gel and purified using the Wizard Genomic DNA Purification Kit (Promega). In order to identify the specific edition, the PCR fragments were Sanger sequenced in Macrogen, Korea. The results were analyzed using Synthego’s ICE (https://www.ice.synthego.com/) online tool^26^.

### 8. Analysis of NK degranulation

To analyze the NK cell degranulation, peripheral blood mononuclear cells (PBMCs) were freshly isolated from buffy coat leukocyte concentrates obtained from anonymous healthy human donors using Ficoll, Histopaque-1077 (Sigma)^27^. Then, NK lymphocytes were isolated from PBMCs by bead magnetic separation. Briefly, the PBMCs were incubated with anti-CD56/biotin antibody (Invitrogen), and then magnetic anti-biotin microbeads were added (Miltenyi). Finally, cells were isolated using a MS-column (Miltenyi) in a miniMACS Separator. After three washes with wash solution (DPBS+ 0.1% albumin + 2mM EDTA), the NK cells were eluted and cultured in RPMI 1640 (Gibco) + 10% FBS. The purification efficacy was analyzed by FC using the following surface markers: anti-CD45/PE antibody (BD Bioscience), anti-CD56/BB515 antibody (BD Bioscience) and anti-CD3/PE antibody (Invitrogen), 1:50 dilution. Once NK cells were isolated, they were incubated with 2 μL rIL-2 (100 U/μL) overnight, to stimulate cellular growth. After 24hs, target cells were co-cultured 1:1 with NK cells in a medium containing 2 μM monensin (eBioscience); cell stimulation cocktail (eBioscience), and anti-CD107a/PE antibody (Invitrogen). After of 4 hs incubation, NKs from all conditions were washed and stained with anti-CD56/BB515 antibody and then analyzed by flow cytometry.

### 9. Statistical analysis

Experimental results are presented as mean ± standard error of the mean (SEM). Statistical significance between groups was analyzed using ANOVA. Residuals fitted normal distribution and homogeneity of variance. Comparisons between means were assessed using Tukey test. For degranulation assay, the paired Student’s t test was performed. Statistical analyses were performed using Infostat Software using 95% confidence intervals.

## Acknowledgements

The authors thank Darío Fernandez Espinosa for his technical assistance.

## Author contributions

Conceptualization: MBP, DTLR, MG, SM.

Formal analysis: MBP, DTLR, LM, MG, SM.

Methodology: MBP, GA, SC, AMM, MAS, MD, IP, LM.

Resources: EC, MG, SM.

Writing-original draft: MBP, DTLR, LM, MG, SG.

Writing-review & editing: MBP, DTLR, ALG, AMI, LM, MG, SG.

## Supplementary information

**Figure S1:**
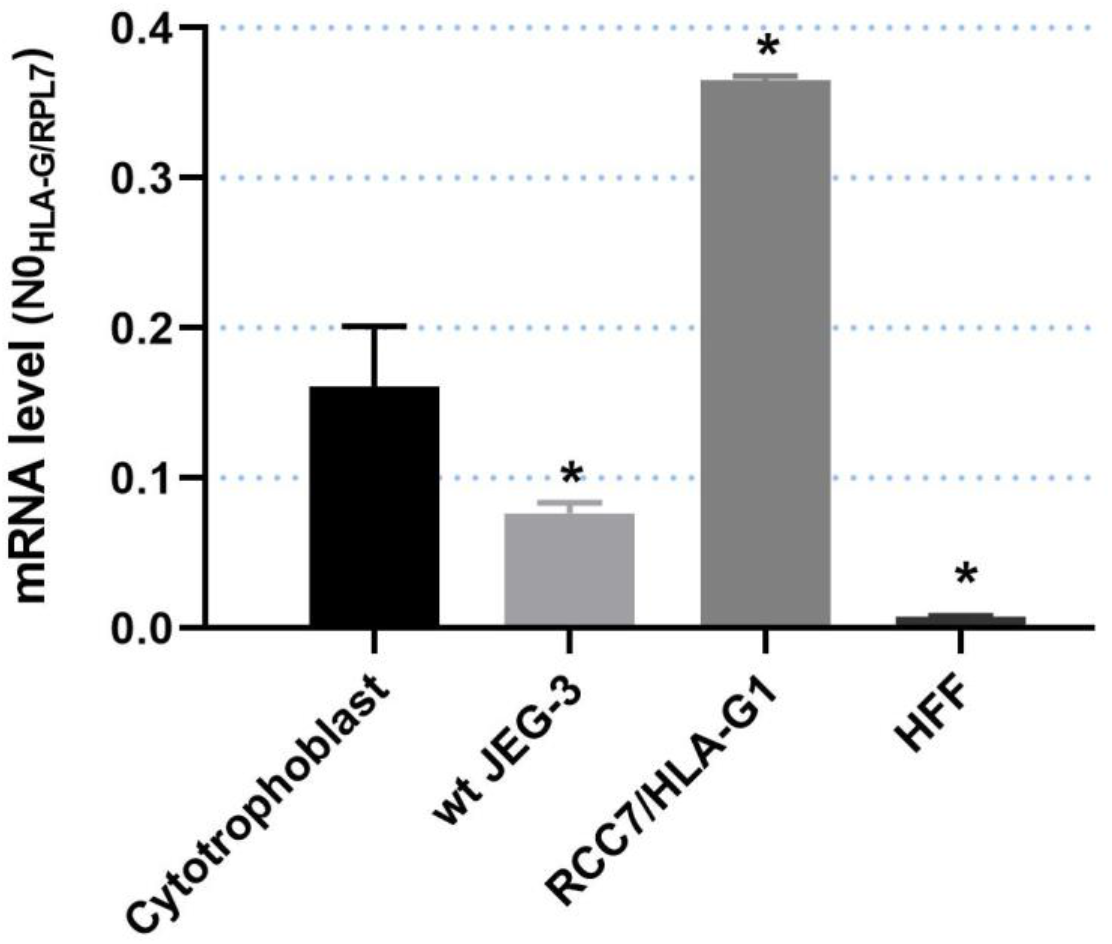
HLA-G expression in JEG-3 and RCC7/HLA-G1 cell lines determined by RT-qPCR. HLA-G expression values were compared to cytotrophoblast cells (extravillous cytotrophoblast extracted from term placenta). HFF: Human fibroblast cells (negative control). JEG-3: choriocarcinoma cell line. RCC7/HLA-G1: renal cell carcinoma cell line expressing HLA-G1. Significant differences are shown with * (p<0.05).

